# Rescuing auditory temporal processing with a novel augmented acoustic environment in a mouse model of congenital SNHL

**DOI:** 10.1101/2020.08.19.256396

**Authors:** Adam C. Dziorny, Luisa L. Scott, Anne E. Luebke, Joseph P. Walton

## Abstract

Congenital sensorineural hearing loss (SNHL) affects thousands of infants each year and results in significant delays in speech and language development. Previous studies have shown that early exposure to a simple augmented acoustic environment (AAE) can limit the effects of progressive SNHL on hearing sensitivity. However, SNHL is also accompanied by “hidden hearing loss” that is not assessed on standard audiological exams, such as reduced temporal processing acuity. To assess whether sound therapy may improve these hidden deficits, a mouse model of congenital SNHL was exposed to simple or temporally complex AAE. Peripheral function and sound sensitivity in auditory midbrain neurons improved following exposure to both types of AAE. However, only exposure to a novel, temporally complex AAE significantly improved a measure of temporal processing acuity, neural gap-in-noise detection in the auditory midbrain. These experiments suggest that targeted sound therapy may improve hearing outcomes for children suffering from congenital SNHL.

## Introduction

Early childhood sensorineural hearing loss (SNHL) is a common neurosensory disability causing significant medical, social and financial hardship. The prevalence of moderate-to-profound SNHL in children (> 40 dB) is roughly 3 in 1,000, with up to 10% have hearing loss that is considered “profound” ^1-4^. There are numerous causes of congenital or acquired sensorineural hearing loss, including genetic factors, infectious diseases, or environmental toxins. Beyond hearing threshold deficits seen in children with SNHL, studies have also shown functional deficits in the development of speech and language processing^5-9^. Impairments in speech perception, which may give rise to these functional deficits, have been associated with restricted encoding of auditory temporal cues ^10^.

Psychoacousticians have used the gap detection paradigm to evaluate temporal processing acuity of sounds for more than 30 years. Gap detection acuity may underlie perceptual boundaries in natural language, such as voiced versus voiceless speech sounds. Minimal gap thresholds (MGT) appear to determine the perceptual boundary in the continuum of voice onset times (VOTs), the intervals between consonant release and the start of vocal cord vibration in consonant-vowel transitions ^11^. Gap detection acuity is also linked to speech recognition abilities ^12^, as well as normal language development ^13, 14^. In animal models, gap detection can be assessed using several different behavioral techniques ^15, 16^, and can also be measured neurophysiologically to assess neural sound encoding. Interestingly, nearly all mammals have similar behavioral MGTs, which are on the order of 2-3 ms; the lowest neural MGTs approximate these behaviorally-assessed MGTs ^17^.

There are several mouse models of congenital SNHL that mimic the different types and progression of hearing loss that occur in humans. The DBA strain, the oldest inbred mouse strain ^18^, contains a mutation in the gene, *Cdh23* ^19^, as well as a nucleotide substitution in *Fscn2* that is the cause of the *ahl8* quantitative trait locus ^20, 21^. This strain shows a rapid, progressive loss of peripheral function beginning at the onset of hearing ^22, 23^, displaying many of the audiometric characteristics found in infants with progressive SNHL^24^. DBA mice have early and rapid loss of outer hair cell (OHC) function in a base to apex progression, as measured by distortion product otoacoustic emission (DPOAE) thresholds ^25^.

Previous studies have shown that when newborn DBA mice are exposed to broadband sounds daily during 12-hour on/off cycles, improvements are seen in peripheral function ^26, 27^, preserving hearing sensitivity and limiting hair cell loss ^28^. The mechanism through which the slowing of the degenerative processes occurs is unknown but perhaps the AAE maintains afferent neuronal input to the central auditory system (CAS) ^29^. An augmented acoustic environment (AAE) exposure also limits neuronal loss in the cochlear nucleus ^28^ and expands the frequency range to which IC neurons are sensitive across the dorsoventral axis compared to non-exposed mice ^30^. When normal-hearing, young adult CBA mice are exposed to AAE no effects, positive or negative, are observed ^31^. Clearly, in mouse models of congenital SNHL, AAE exposure shows promise in ameliorating the effects of rapid, progressive SNHL on loss of hearing sensitivity. However, whether AAE exposure can ameliorate other aspects of auditory dysfunction associated with SNHL has yet to be studied. The goal of the current study was to test the hypothesis that exposure to targeted AAE having complex temporal sound features would improve neural correlates of gap encoding in the CAS.

## Materials and Methods

### Animal Model

The DBA/2J mouse strain has served as a mouse model of early-onset severe hearing loss for over 4 decades ^22, 23, 32^. Founder breeding pairs (JAX 000671) were obtained from Jackson Labs (Bar Harbor, ME) and our colony was maintained in micro-isolator facilities in the institutional vivarium. Mice were housed in rodent micro-isolator cages and provided *ad lib* food and water. Lights in the room were on a 12-hr light/dark cycle. Cages were changed at least weekly and mice were monitored for signs of distress by trained vivarium technicians. Breeder pairs were kept separate from experimental mice; only nulliparous mice were used for experiments. Both control and exposed pups were weaned into gender-separated cages between ages postnatal day (P) P21 and P28. All mice in this study were between 1^st^ and 4^th^ generation Jackson Labs breeder mice offspring. All procedures were approved by the University of Rochester’s Committee on Animal Resources and in strict accordance with the National Institutes of Health Guide for the Care and Use of Animals.

### AAE Exposure

Mice were exposed using the same amplifier and sound source as described previously ^26^ (generously provided by Dr. Jeremy Turner). Cages were housed inside a sound-attenuating chamber (∼ 3 feet wide × 2 feet deep × 5 feet high) covered with anechoic foam, and the booth itself was housed in a 2-way vivarium room. Stimulus presentation was calibrated *in situ* to 70 dB SPL using a Quest 1900 Sound Level Meter and an ACO Pacific ¼” free-field microphone (Figure 1A). The spectrum of regular AAE (R-AAE) exposure was recorded using an HP/Agilent 35665A Spectrum Analyzer (Hewlett Packard). The analysis revealed a wide-band noise ± 6 dB from 4 – 20 kHz (Figure 1B). Ambient noise levels were between 39 – 45 dB SPL in this frequency range. Our novel temporal AAE (T-AAE) stimulus was generated from a subsection of the wav file containing our original stimulus (Figure 1C), with additional silent gaps inserted within the noise bursts, as follows. Random gap durations of 0, 1, 2, 4, 8 or 16 ms were inserted into the wave vector 100 ms into each 200-ms noise burst, and the remaining noise burst was shortened by the same gap duration to preserve the 40% duty cycle. The resulting wave vector was saved and utilized for temporal AAE stimulus presentation (Figure 1C).

**Figure 1.**
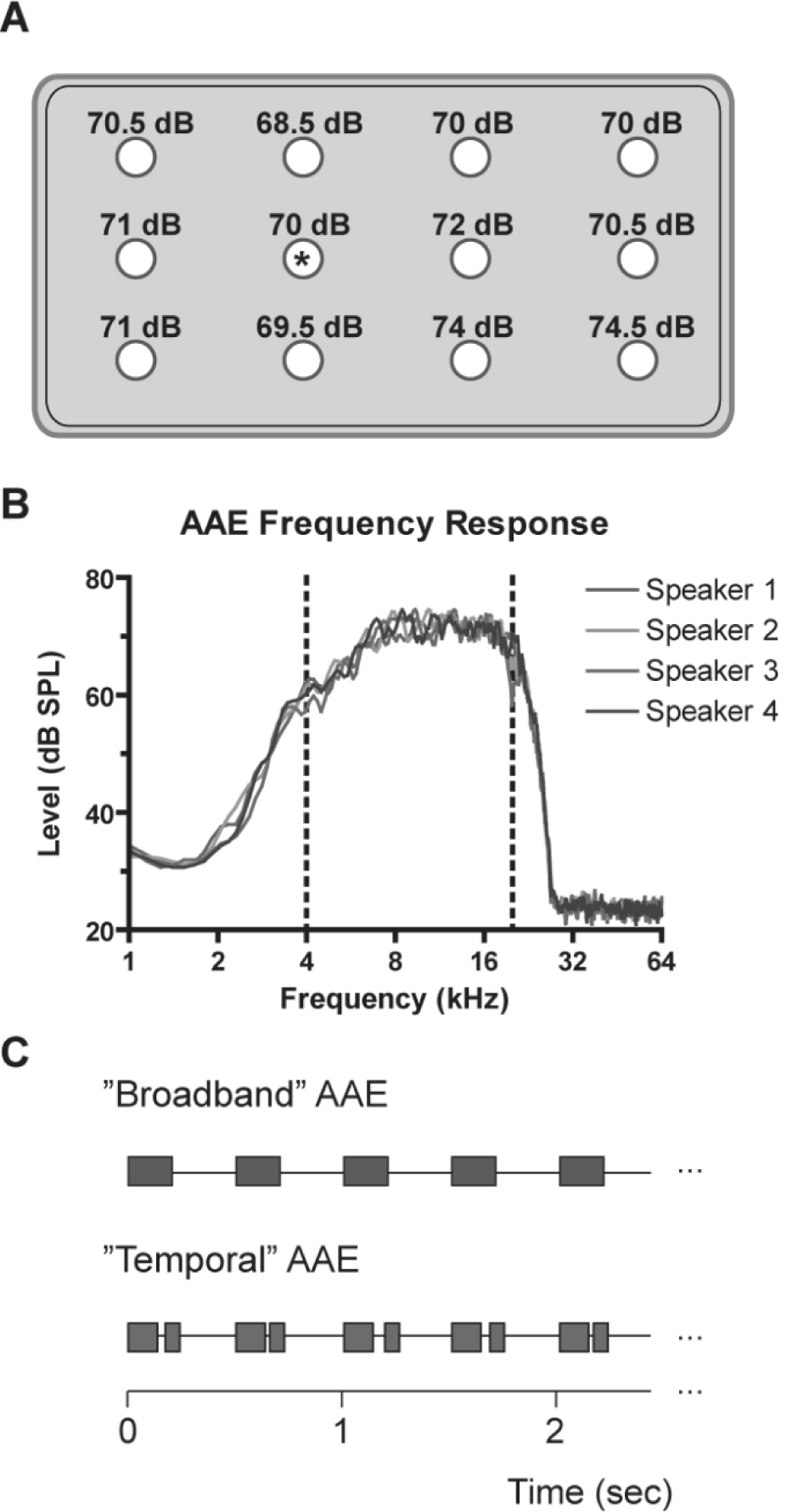
Exposure calibration, stimulus spectrum and temporal pattern. **A**, Sound levels recorded at different points in the cage (circles, spaced approx. 3.5” apart) in response to calibrated AAE stimulus. Asterisk denotes hole calibrated to 70 dB SPL. **B**, The frequency response spectrum of the AAE stimulus, presented through each speaker used during exposure, demonstrates a flat region (± 6 dB) from 4 – 20 kHz (indicated with dashed lines). The spectrum was recorded with a ½” ACO Pacific microphone on a Quest 1900 sound level meter output to an HP/Agilent 35665A spectrum analyzer. Each waveform consisted of 30 averages. **C**, Schematic of regular and temporal augmented acoustic environment exposure. Each exposure was presented twice per second, 200 ms per burst, for 12 hrs / day. Temporal AAE stimulus had a silent gap (either 0, 1, 2, 4, 8 or 16 ms in duration) inserted after the first 100 ms.

AAE exposure began just after birth, before the onset of hearing at around P10 ^33^ matching previous studies in the DBA model ^26, 28, 31^. All mice in this study were tested at 30 days after birth, following at least 18 days of AAE exposure. This time point was chosen because control mice of this strain already show significant hearing loss by this time ^22, 23, 26^.

### Peripheral Auditory Assessment

Auditory brainstem responses (ABRs) were recorded using BioSigRP software (version 4.4.1, Tucker-Davis Technologies, Alachua, FL) interfacing with Tucker-Davis Technologies (TDT) System III hardware. ABR waveforms were recorded in response to tone bursts of 5 ms duration, shaped by a Blackman window. The frequencies tested were 3, 6, 12, 16, 20, 24, 32 and 36 kHz for each animal. Stimuli were presented at a rate of 25 / second, with 150 averages per waveform, with replication. Artifact rejection was enabled with a threshold of 7 μV. Each frequency was presented beginning at a level of 80 dB SPL down to 20 dB below threshold, in 5-dB increments. The recorded waveforms were amplified (x10,000), filtered (0.3 – 10 kHz) and digitized. No mice (control or experimental) responded to test frequencies >24 kHz, and therefore these frequencies were omitted from the ABR analysis.

Distortion product otoacoustic emissions (DPOAE) amplitudes were measured using custom MATLAB (Mathworks) software interfacing with TDT System III hardware, calibrated similar to the ABR acquisition hardware. The speaker transduction tube / ER10B microphone apparatus was lowered into the ear canal of the anesthetized mouse using a micromanipulator under microscopic examination. Two separate placements & recordings were completed on each mouse; if the results differed, a third placement and recording was completed. The pair of matching results was averaged during analysis. DPOAE amplitudes were recorded in response to two simultaneous pure-tone bursts (f_1_ and f_2_) at different frequencies, related with the following ratio: f_2_ / f_1_ = 1.25. The lower-frequency tone (f_1_) was presented at 65 dB and the higher-frequency tone (f_2_) was presented at 50 dB. The geometric mean presentation frequencies were from 5.6 – 20.5 kHz. Amplitudes were transformed to the frequency domain and the cubic distortion product (2f_1_ – f_2_) and surrounding noise values were measured. DPOAE responses could not be distinguished from the noise floor above 22 kHz in any group, and these frequencies were not analyzed.

### Auditory Midbrain Neurophysiology

We recorded neuronal activity in the inferior colliculus (IC) using a 16-channel vertically-oriented electrode (a1×16-3mm-100-177μm^2^, NeuroNexus Technologies) with 100 μm spacing between pads and impedances of 1 – 3 MΩ. Electrodes were positioned over the craniotomy and advanced ventrally by a micromanipulator. The electrical output was amplified, filtered and digitized at 25 kHz in a 1.25 ms time window. Neural activity was automatically determined using a 3:1 SNR.

Stimuli were generated using DSP software (OpenEX and RPVDS, TDT) and presented through TDT System III hardware to an electrostatic speaker (TDT ES1). The speaker was located at a 60° azimuth contralateral to the recording site. Stimulus presentation was controlled by custom MATLAB routines interfacing through OpenEx interfaced to an RX6. First, search stimuli were presented to locate responsive units and identify events. These stimuli were band-limited noise bursts (3 – 50 kHz) presented at 70 dB SPL at a rate of 2 / second. Second, tone burst stimuli (25-ms in duration, 10 / sec) were presented to measure frequency response areas (FRA). The range of frequencies used in this study was 2 – 64 kHz (500 Hz increments) and the range of levels was 0 – 85 dB SPL (5 dB increments). Each frequency-level pair was presented five times, with the entire set randomized prior to presentation. Third, to assess gap-in-noise encoding noise bursts of 100 ms (noise burst 1, NB1) and 50 ms (noise burst 2, NB2) were delivered at a rate of 2 / sec. The level was fixed at the start of each run to 80, 70, or 60 dB SPL, as these intensities were predicted to be >20 dB higher than the noise threshold for individual units. Silent gaps in the noise burst were inserted (0.25 msec rise-fall) after the first 100 ms of the first noise burst (NB1), with the gap duration being one of the following: 0, 1, 2, 4, 8, 16, 32, 64, or 96 ms. Continuous background noise (CBN) was used to further test the benefits of AAE exposure. CBN (3-50 kHz) could be applied to a gap series at a fixed level (+6 dB SNR) whereby the silent gap would also be filled with this continuous background noise at a level of 6 dB below the noise carrier. Each gap duration was repeated 50 times, for a total of 500 repetitions (10 gap durations × 50 repetitions per duration).

### Spike Sorting and Response Measures

Spike waveforms were processed in MATLAB® using the TDT OpenDeveloper ActiveX controls and passed to AutoClass C v3.3.4, an unsupervised Bayesian classification system that seeks a maximum posterior probability classification, developed at the NASA Ames Research Center ^34^. AutoClass scans the dataset of voltage–time waveforms according to custom specified spike parameters to produce the best-fit classifications of the data, which may include distinct single- and multi-unit events, as well as noise. To discriminate the signal from noise, the variance of the background noise was estimated as the quartile range of the first five digitization points of the spike waveform, as these are recorded prior to the threshold-crossing event. To avoid overloading AutoClass with excessive noise, which leads to over-classification, this noise measure was used to screen the event waveform data such that only voltage points with absolute values greater than this noise floor were presented for use in the classification. Once the classes had been determined in each channel of data, they were visualized within a custom MATLAB® program and assigned to multi-unit, single-unit, or noise classes. Event classes which were categorized as noise were subsequently discarded, and units with distinct biphasic waveforms and good SNR were classified as single-units. As most channels recorded information elicited from the spiking of two or more neurons, all recordings units in this paper were considered to be multi-unit activity ^35^. Nonetheless, there was no observation of any consistent differences in the eFRAs between single units and multi-unit clusters.

Data analysis was performed as previously described ^36^. Frequency response areas (FRAs) were displayed in a custom MATLAB GUI and analyzed with a multi-step procedure using custom software. Frequency receptive fields (FRAs) were then used to determine the best frequency (BF), the frequency with the lowest intensity of driven activity, and tuning sharpness. For units with sound driven activity, assessed within the FRA, neural responses to gap-in-noise were visualized using a custom Matlab GUI, and minimum gap thresholds (MGTs) were determined using previously-published methods ^37, 38^. Only gap-responsive units (with MGT ≤ 96 ms) for each stimulus condition were included in subsequent analysis of gap detection. Additionally, the analysis focuses on phasic units because tonic units recorded in continuous background noise demonstrated post-excitatory suppression. Due to this post-excitatory suppression, the quiet window responses of tonic units were not strictly a result of the embedded silent gap, making MGT determination highly variable and unreliable.

### Statistical Analysis

Table 1 reports the number of mice that underwent ABR, DPOAE, and IC recordings, and the number of IC units included in each measure of neural sound processing. Mice with DPOAE recordings were a subset of mice with ABR recordings; however, mice that underwent IC recordings did not always have peripheral assessment. Auditory processing measures are reported as mean ± standard error of the mean and statistical comparisons were made using GraphPad Prism v6.0. The Student’s t-test compared differences between two groups, while Analysis of Variance (ANOVA) with Bonferroni *post-hoc* testing compared the effect of one or more variables. The Chi Square or Fischer’s Exact test was used to examine differences between observed and expected counts. Significance was set at *p* < 0.05.

**Table 1.** Counts of animals tested and units isolated. For peripheral measures, counts are given in terms of the number of animals tested. For central auditory recording, counts are listed in terms of the number of animals as well as the number of units recorded from these animals. Percent values are listed as percent of total units recorded for each exposure type.

|  | **Control** | **Regular** | **Temporal** |
| --- | --- | --- | --- |
|  |  | **AAE** | **AAE** |
| *Periphery (# Mice)* |  |  |  |
| ABRs | 20 | 16 | 15 |
| DPOAEs | 13 | 12 | 8 |
| *Auditory Midbrain* |  |  |  |
| # Mice | 11 | 11 | 10 |
| Total # Units Recorded | 825 | 932 | 904 |
| Frequency Response Areas (%) | 574 (69.6) | 702 (75.3) | 629 (69.6) |
| 80 dB Gap-Responsive (%) | 507 (61.5) | 595 (63.8) | 567 (62.7) |
| 80 dB Phasic Units (%) | 242 (29.3) | 192 (20.6) | 267 (29.5) |
| 80 dB Gap-In-Noise Responsive | 173 (21.0) | 134 (14.4) | 210 (23.2) |
| 70 dB Phasic Units (%) | 204 (24.7) | 184 (19.7) | 234 (25.9) |
| 60 dB Phasic Units (%) | 96 (11.6) | 170 (18.2) | 188 (20.8) |

## Results

### Peripheral Auditory Assessment

ABRs were recorded to determine the effect of early AAE exposure beginning at the onset of hearing. ABR thresholds were assessed as a function of Exposure and Frequency (Figure 2A, D). A two-way ANOVA demonstrated significant effects of Exposure (*F* = 23.46, *p* < 0.001), Frequency (*F* = 121.10, *p* < 0.001) and Exposure × Frequency (*F* = 5.71, *p* < 0.001) with post AAE thresholds from exposed mice being significantly improved as compared to control mice. Post-hoc analysis showed significant improvement in ABR thresholds at 12 and 16 kHz following either type of AAE exposure, and no significant differences between the two types of AAE exposure. The magnitude of the difference in ABR thresholds for control and AAE-exposed mice approached 30 dB at 16 kHz (Control vs. Regular AAE: 29 dB; Control vs. Temporal AAE: 26 dB), a frequency in the range of the best hearing for CBA mice. While the difference did not reach statistical significance, at 24 kHz we encountered very few mice from the Control group that had observable responses at 80 dB (3 / 20, or 15%), when compared to the Regular AAE (6 / 16, or 38%) or Temporal AAE groups (7 / 15, or 46%). Together, these findings indicate that ABR thresholds improved following exposure to both types of AAE (Figure 2A, D), replicating the findings of Turner and Willott (1998). Moreover, the frequency range that showed the most improvement in ABR thresholds was within the region of maximal energy for the AAE exposure spectrum and with the frequency region of best hearing sensitivity.

**Figure 2.**
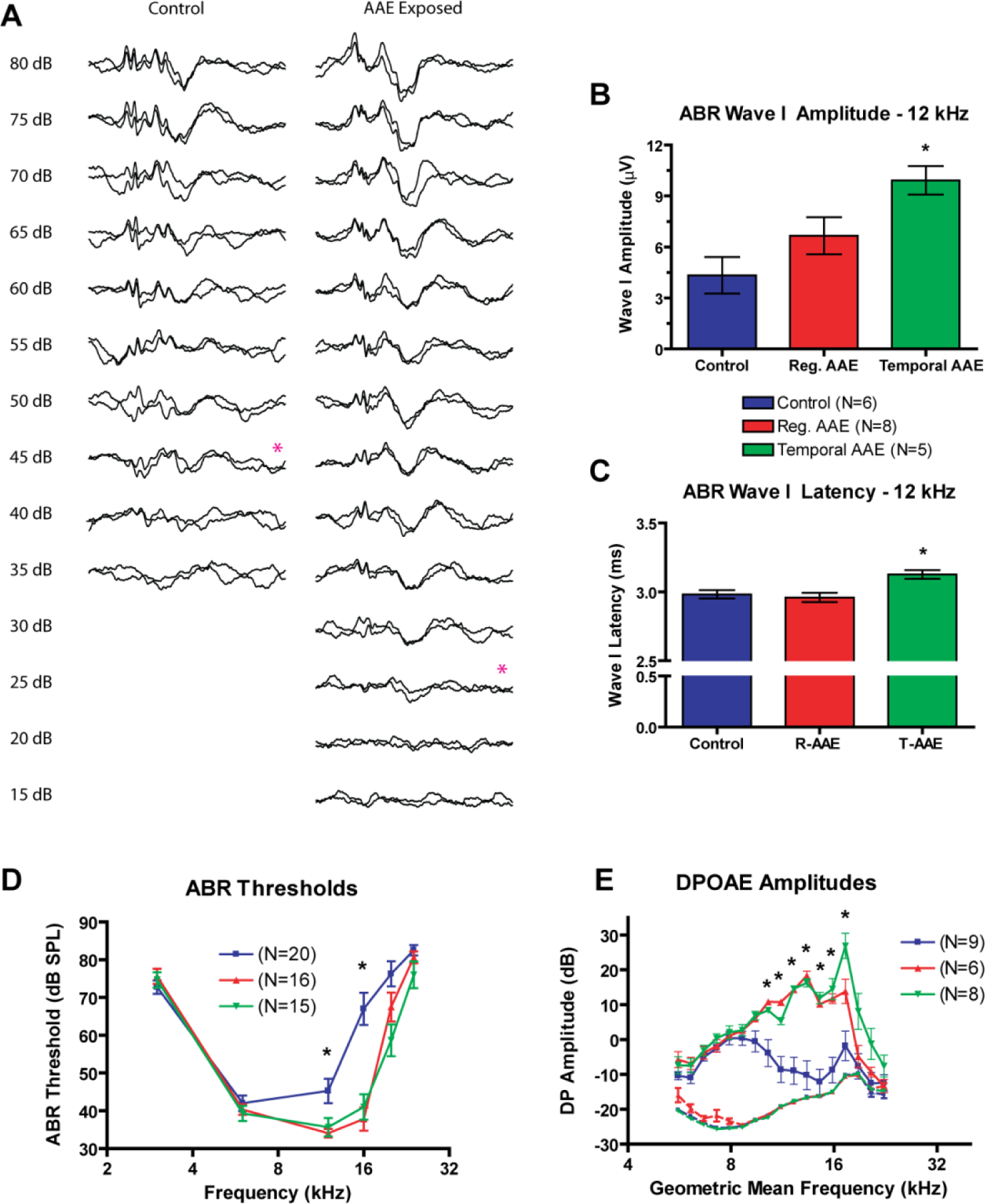
AAE exposure improves peripheral function at P30. **A**, Representative ABR waveforms from a control and AAE-exposed animal at 12 kHz show similar suprathreshold morphology, with an elevated threshold in the control animal. **B**, ABR wave I amplitudes were increased in both exposure types compared to controls (Control *[blue]*: 4.3 ± 1.1 μV; Reg. AAE *[red]*: 6.7 ± 1.1 μV; Temporal AAE *[green]*: 9.9 ± 0.8 μV), though only responses from temporal AAE exposure reached significance (*p* < 0.05). **C**, ABR wave I latencies were similar in magnitude (Control: 2.98 ± 0.03 ms; Reg. AAE: 2.96 ± 0.03 ms; Temporal AAE: 3.13 ± 0.03 ms). Temporal AAE-exposed mice had significantly longer latency compared to control mice (p < 0.05) and regular AAE mice (*p* < 0.01). **D**, ABR thresholds were significantly decreased at 12 & 16 kHz following exposure to both types of AAE (Regular = *red*, Temporal = *green*) compared to controls (*blue, p* < 0.001). At the frequency of greatest differences (16 kHz), this difference approaches 30 dB. No responses were noted above 24 kHz in any group. **E**, DPOAE amplitudes were increased following exposure to both types of AAE (Regular = *red*, Temporal = *green*) compared to controls (*blue*), with larger increases seen at select frequencies following temporal AAE exposure. Between 10 and 17 kHz, both types of AAE exposure significantly increased amplitudes (mean increase: 21.2 ± 1.4 dB, *p* < 0.01). Additionally, temporal AAE exposure resulted in a 13 dB amplitude increase over regular AAE exposed mice at two test frequencies (17.27 & 18.84 kHz). Amplitudes could not be distinguished from the noise floor above 22 kHz in any group.

The amplitude and latency of wave I in the ABR waveform was measured in response to a 12 kHz tone presented at 80 dB SPL (Figure 2B, C). Wave I amplitudes (Figure 2B) demonstrated a significant effect of Exposure (*F* = 5.94, *p* = 0.0118). Post-hoc group comparisons reveal wave I amplitudes were significantly greater for temporal AAE-exposed mice than controls (4.3 ± 1.1 μV compared to 9.9 ± 0.8 μV, *p* < 0.05). Mean wave I amplitude was also greater for regular AAE-exposed mice (6.7 ± 1.1 μV) than controls, though the difference did not reach significance. Although the mean wave I latencies were physiologically similar across groups, differing by only 170 µsec (Control: 2.98 ± 0.03 ms; Regular AAE: 2.96 ± 0.03 ms; Temporal AAE: 3.13 ± 0.03 ms; Figure 2C) the one-way ANOVA demonstrated a significant effect of Exposure (*F* = 6.81, *p* = 0.007). Together these findings indicate the temporal AAE exposure had a more substantive effect on wave I ABR measures of cochlear sound processing than regular AAE exposure.

DPOAE amplitudes were measured in mice following exposure to regular or temporal AAE versus control environments to assess outer hair cell function (Figure 2E). A two-way ANOVA found significant effects for Exposure (*F* = 109.00, *p* < 0.001), Frequency (*F* = 17.98, *p* < 0.001) and Exposure × Frequency (*F* = 5.25, *p* < 0.001). Post-hoc group comparisons revealed that DPOAE amplitudes increased at geometric mean frequencies between 10 – 17 kHz for both groups of AAE exposure relative to control (mean increase for pooled AAE-exposure responses: 21.2 ± 1.4 dB, *p* < 0.05). Temporal AAE exposure resulted in an even larger impact on DPOAE amplitudes than regular AAE exposure. Post-hoc group comparisons revealed that DPOAEs elicited by 17.27 and 18.84 kHz tones were significantly larger (by 13 dB) in temporal AAE-exposed animals compared to regular AAE-exposed animals (*p* < 0.05). Thus, similar to the ABR analysis, the DPOAE analysis showed the greatest impact of AAE exposure on cochlear function for stimuli in the frequency region with maximal AAE energy. Additionally, Temporal AAE exposure was moderately more effective at improving DPOAE measures of cochlear function.

### Central Auditory Function

To determine whether AAE exposure can influence neural markers of spectral or temporal auditory processing acuity, we measured the response of IC neurons to sound stimuli in vivo. To our knowledge this is the first description of the effects of AAE on neural coding of complex sounds in the CAS. Exposure to AAE altered the frequency response properties for IC units when compared to units from control, unexposed animals (Figure 3A-D). One-way ANOVAs demonstrated that exposure to AAE resulted in a significant upward shift in BF (*F* = 43.16, *p* < 0.001) and a significant improvement in minimum threshold (*F* = 15.46, *p* < 0.001) of neurons from AAE groups. Group comparisons revealed that both types of AAE exposure significantly increased the upward frequency boundary of BFs relative to Control values (Reg. AAE: 14.2 ± 0.2 kHz; Temporal AAE: 15.1 ± 0.2 kHz; Control: 12.4 ± 0.2 kHz; *p* < 0.001), with no further differences between regular and temporal AAE exposure groups (Figure 3B). Likewise, both types of AAE exposure significantly improved minimal response threshold, with no further effect of AAE exposure group (Figure 3C). Finally, exposure to either type of AAE sharpened tuning of the FRAs, assessed by measuring Q-values for the FRA between 10 and 40 dB above threshold. One-way ANOVAs demonstrated a significant effect of Exposure for Q_10_ through Q_40_ values (Q_10_: *F* = 28.81; Q_20_: *F* = 43.46; Q_30_: *F* = 34.09; Q_40_: *F* = 48.58; *p* < 0.001). A higher Q-value indicates sharper tuning, and group comparisons showed that Q_10_ through Q_40_ values following either type of AAE exposure were higher than control values (Q_10_ shown in Figure 3D). The mean magnitude of the difference between control and AAE exposure groups at Q_10_ is approximately 1, which at a BF of 12 kHz equates to a bandwidth difference of about 1 kHz (or 25% of the Control group mean). No other significant differences were found with respect to frequency receptive fields between exposure types. These findings indicate that in AAE-exposed mice, IC neurons had lower thresholds and were more responsive to higher frequencies, but were also more narrowly tuned. The type of AAE exposure did not influence these improvements in spectral acuity.

**Figure 3.**
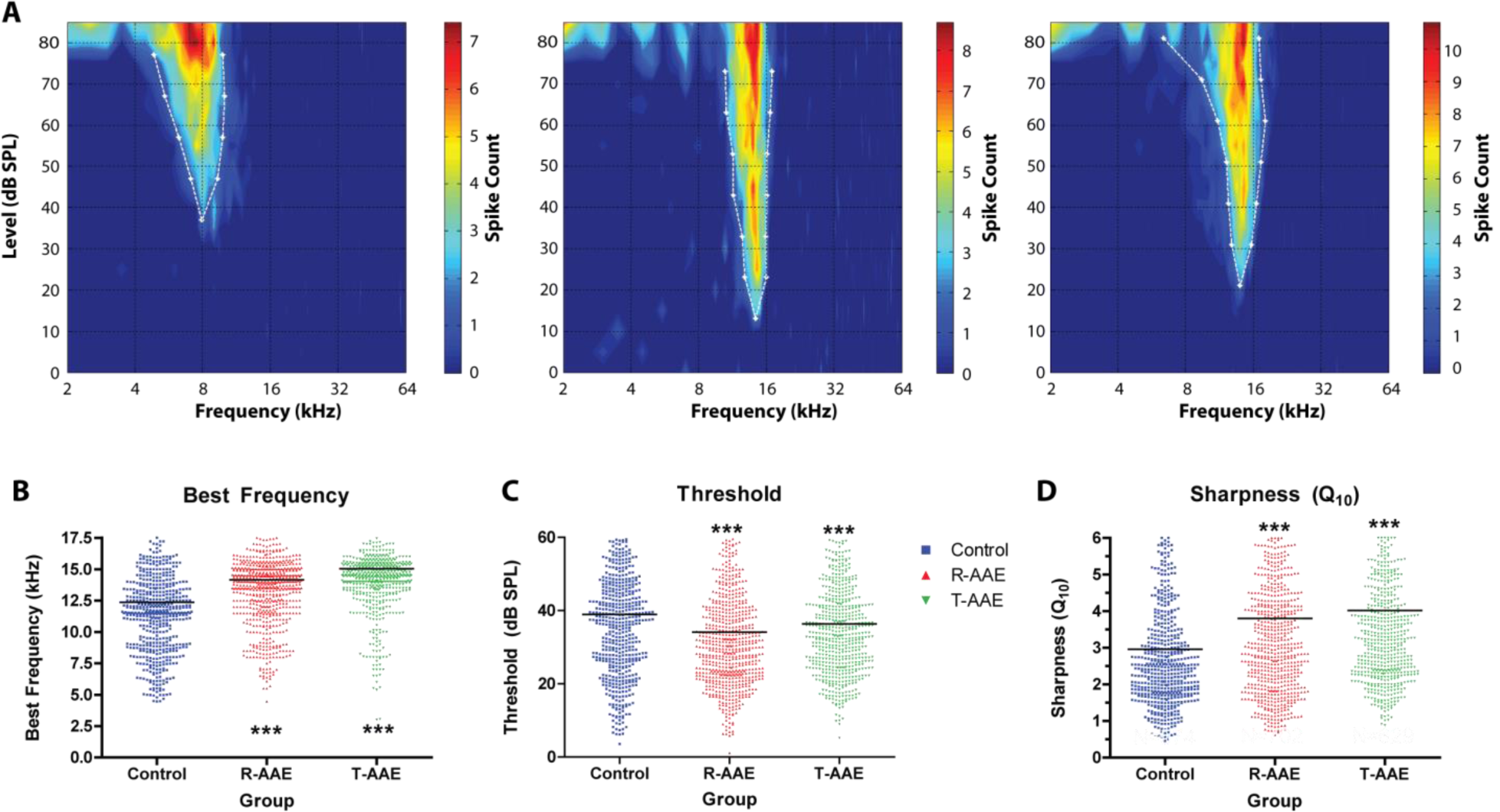
Exposure to AAE changes the frequency responses of IC units. **A**, Example frequency response areas (FRA) are shown from representative control, regular AAE-exposed and temporal AAE-exposed animals. Color-mapped counts indicate the number of spikes per frequency-level pair, with the legend shown on the right. Best frequency (BF) was automatically identified as the frequency with the lowest intensity of drive (minimum threshold, or MT). **B**, Mean best frequencies were significantly increased compared to controls following either type of AAE exposure (Reg. AAE: 14.2 ± 0.2 kHz, Temporal AAE: 15.1 ± 0.2 kHz, Control: 12.4 ± 0.2 kHz; *p* < 0.001). No significant difference was seen between AAE exposure types. **C**, Mean unit thresholds were significantly decreased following either type of AAE exposure (Reg. AAE: 34.2 ± 0.6 dB, Temporal AAE: 35.9 ± 0.5 dB, Control: 38.9 ± 0.7 dB; *p* < 0.01). Again no significant difference was seen between AAE exposure types. **D**, Tuning sharpness was improved with both types of AAE exposure. Q-values computed at 10 dB as well as 20 dB, 30 dB and 40 dB above threshold (data not shown) were significantly increased compared to controls (one-way ANOVA with post-hoc testing, *p* < 0.001 at all levels). No significant differences were seen between the two types of AAE exposure. Graphs in B, C and D were vertically scaled to demonstrate differences in the mean, and thus some data points above the maximum vertical axis value are not shown.

Temporal processing acuity was assessed via MGTs, a measure of neural coding of silent gaps embedded in noise. Representative post-stimulus time histograms (PSTHs) of single phasic units in response to different gap durations are shown in Figure 4. Minimum gap thresholds (MGTs) were computed for all phasic units, and units were included in each of the following analyses if the gap threshold for the condition was ≤ 96 ms (Figure 5). For gaps embedded in 80-dB noise carriers, one-way ANOVA demonstrated a significant effect of Exposure (*F* = 15.43, *p* < 0.001; Figure 5A). Both types of AAE exposure shortened MGTs relative to controls. The mean magnitude of improvement for the regular AAE exposure group was 4.89 ms (33%), and for the temporal AAE exposure group was 6.58 ms (44%). For 70-dB carriers, one-way ANOVA demonstrated a significant effect of Exposure (*F* = 12.49, *p* < 0.001). Again, both types of AAE exposure improved MGTs compared to controls, with greater average improvement seen by mice exposed to temporal AAE (8.10 ms) versus those exposed to regular AAE (5.11 ms). Post-hoc comparison did not show a significant difference between groups exposed to regular versus temporal AAE. For silent gaps embedded in 60-dB SPL carriers, one-way ANOVA demonstrated an effect of Exposure on MGT (*F* = 7.31, *p* < 0.001). Group comparisons showed that mice exposed to temporal AAE had significantly shorter MGTs compared to mice exposed to regular AAE (temporal AAE vs. regular AAE: 15.9 ± 1.2 ms vs. 23.3 ± 1.6 ms, p < 0.01). Control mice also had shorter MGTs compared to mice exposed to regular AAE, though these differences did not reach significance (control vs. regular AAE: 18.6 ± 1.7 ms vs. 23.3 ± 1.6 ms). However, the number of units with detectable minimal gap thresholds was also substantially lower for control mice (96 units) than for either regular (170 units) or temporal (188 units) AAE-exposed mice (*χ*^*2*^(2) = 118.08, *p* < 0.001). In contrast with phasic units, tonic units showed no significant effects of AAE exposure on responses to gap stimuli or MGTs (data not shown). Overall, these findings indicate that gap detection generally improved in phasic units following exposure to both types of AAE, with a trend towards greater improvement seen following exposure to our novel temporal AAE.

**Figure 4.**
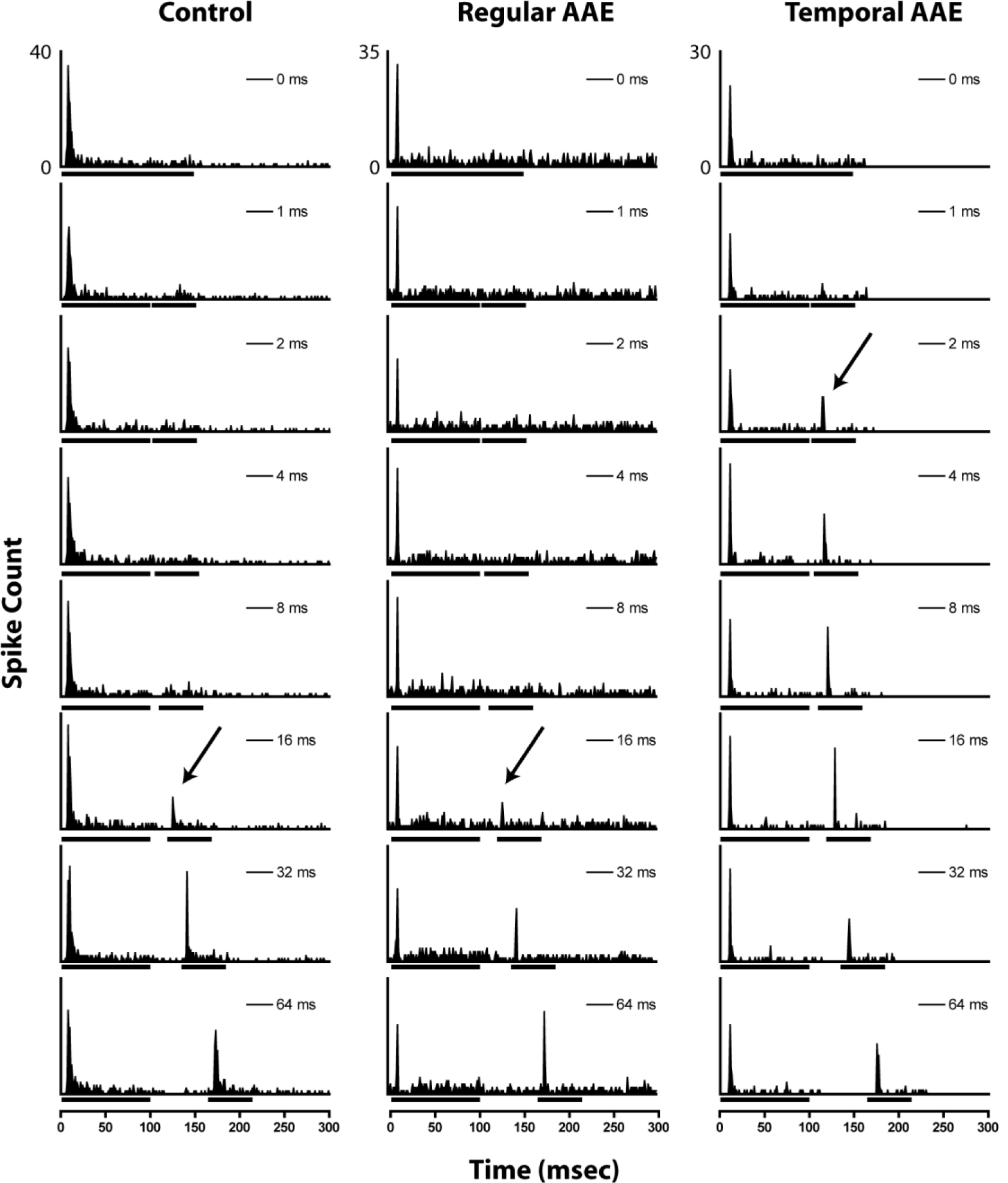
Representative examples of a neural correlate of gap encoding by phasic units from unexposed (*left*), regular AAE-exposed (*center*), and temporal AAE-exposed (*right*) mice. Post stimulus time histograms show spike counts summed over 50 presentation of a gap-in-noise paradigm using a carrier level of 80 dB SPL with gap duration shown in each PSTH. Bars under the x-axis denote noise-burst duration marking the silent gap. Arrows denote the automatically-calculated MGT for each unit.

**Figure 5.**
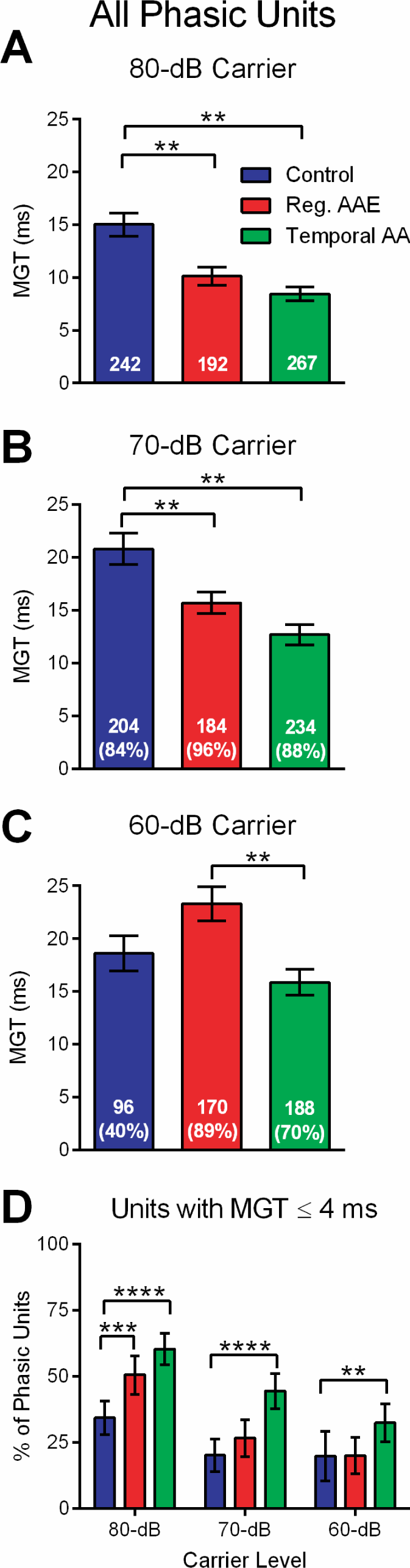
Exposure to both types of AAE improve mean gap thresholds in phasic units. Mean gap thresholds (MGTs) were computed across each group, for each noise carrier level (80, 70 & 60 dB). Total number of units in each group are shown at the bottom of the bar. **A**, At the 80-dB carrier level exposure to both types of AAE resulted in significantly shorter mean MGT (Control: 15.0 ± 1.1 ms, Reg. AAE: 10.1 ± 0.9, Temporal AAE: 8.4 ± 0.7 ms). **B**, At the 70-dB carrier level, exposure to both types of AAE again significantly shorten mean MGT, with greater improvement seen in the temporal AAE exposure group (Control: 20.8 ± 1.5 ms, Reg. AAE: 15.7 ± 1.0, Temporal AAE: 12.7 ± 1.0 ms). **C**, At the 60-dB carrier level the mean MGT from the temporal AAE group was significantly shorter than from the regular AAE group (15.9 ± 1.2 ms vs 23.3 ± 1.6 ms, *p* < 0.01). Additionally, the mean MGT for control mice was significantly shorter than from the regular AAE group (18.6 ± 1.7 ms vs 23.3 ± 1.6 ms) but the number of responsive units was much less. Sample size is shown inside the bar, with the percent equal to the percent of all phasic responsive units to an 80-dB carrier. **D**, A significantly greater number of phasic units had MGTs ≤ 4 msec in mice exposed to temporal AAE, followed by those exposed to regular AAE, for all carrier levels, 80, 70 and 60 dB SPL (** denotes p<0.05, ***=p<0.001, ****=p<0.0001 by Fischer exact test). The fraction of phasic units is displayed with ± 95% CI.

Gap detection is more challenging in background noise, and may also be a key marker of speech recognition difficulties in background noise ^39, 40^. To determine whether this measure of temporal acuity improves following AAE exposure, only units that responded to gap stimuli (MGTs ≤ 96 ms) presented in continuous background noise (CBN) were included in the analysis (see Table 1 and Figure 6A, B). This subpopulation also showed improvement in MGTs for gap stimuli presented in quiet after either AAE exposure (One-way ANOVA: *F* = 18.16, *p* < 0.001; post-hoc regular and temporal AAE MGT < control, *p* < 0.001; compare Figure 6A with Figure 5A). A one-way ANOVA also showed a significant effect of Exposure on MGTs when stimuli were delivered in the presence of +6 dB SNR continuous background noise (*F* = 5.39, *p* = 0.005; Figure 6B). Exposure to temporal AAE significantly shortened MGTs compared to controls (12.7 ± 1.0 ms vs. 17.9 ± 1.2 ms, *p* < 0.01), while exposure to regular AAE only trended towards shorter MGTs (14.6 ± 1.2 ms vs. 17.9 ± 1.2 ms, *p* > 0.05). These data indicate that early temporal AAE exposure improves gap detection in the presence of background noise for phasic units.

**Figure 6.**
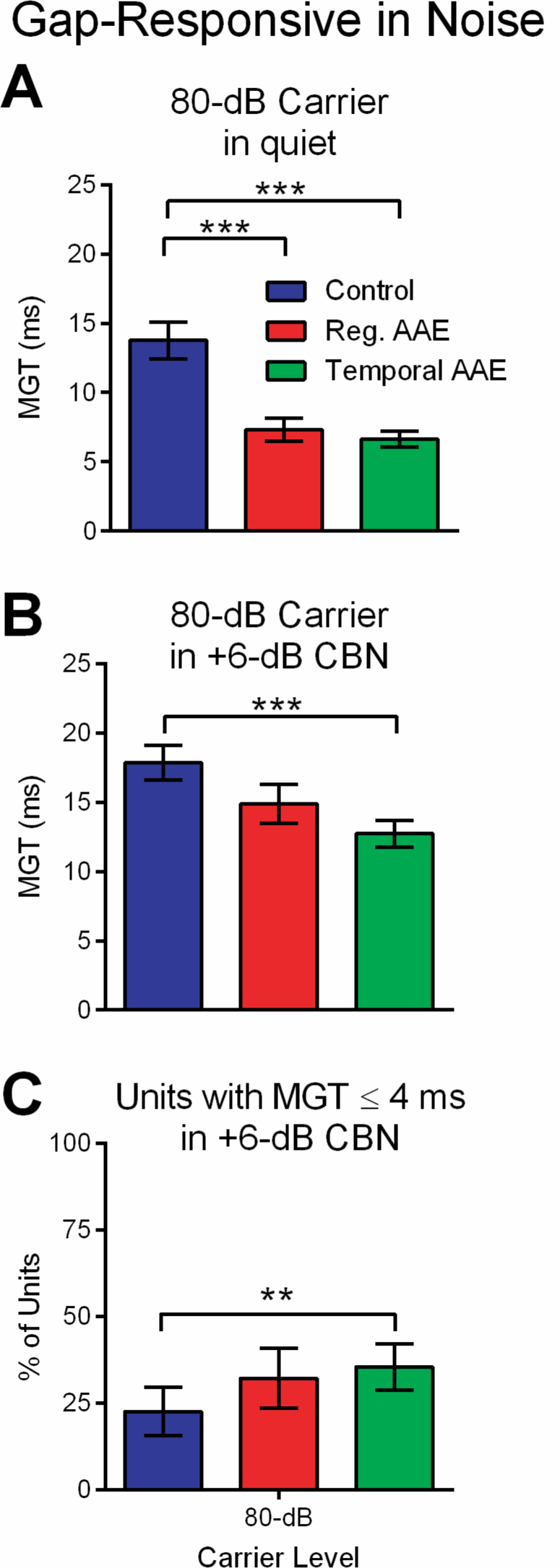
Exposure to temporal AAE preserves gap thresholds in the presence of continuous background noise (CBN). Only a subset of phasic units was responsive in background noise. **A**, Shown are the MGTs measured in response to an 80-dB carrier without CBN from only the units responsive in background noise. These results match those seen in Figure 5A where at 80-dB both types of AAE exposure resulted in significantly shorter MGT. **B**, The same units as in A, measured in response to an 80-dB carrier with +6 dB SNR CBN (background noise at 74 dB). Only exposure to temporal AAE resulted in significantly shorter gap thresholds compared to controls (12.7 ± 1.0 ms vs 17.9 ± 1.2 ms, *p* < 0.01). Sample size is given in each bar, with percent shown as a percent of all phasic responders at 80-dB. **C**, The percent of all responsive phasic units with MGTs ≤ 4 ms (sensitive responders) was increased in the temporal AAE group at all levels (** denotes p<0.05 by Fischer exact test. The error bars show fraction of phasic units with ± 95% CI.

As our previous work suggests that behavioral gap thresholds are more strongly influenced by midbrain units with the shortest gap thresholds ^17^, we computed the percent of responding units with gap thresholds ≤ 4 ms (Figure 6C). All phasic units with MGT ≤ 96 ms (those shown in Figure 5A-C) were included for this analysis. Temporal AAE-exposed mice had the greatest percent of phasic units that had short gap thresholds, regardless of carrier intensity. Thus, early temporal AAE strengthens encoding of short gap durations.

## Discussion

In the present study, we have shown that exposure to a temporally-complex broadband AAE can modulate multiple aspects of peripheral and central auditory function in a mouse model of severe congenital SNHL. Early exposure to a novel, temporally-enriched broadband noise stimulus, starting before hearing onset, improved ABR thresholds, wave I ABR amplitudes and DPOAE amplitudes relative to normally-raised mice. The frequency range that showed the most improvement in cochlear function was within the region of maximal energy for the AAE exposure spectrum. Improvements in sensitivity and spectral encoding were also present in the CAS. Recordings in the auditory midbrain showed lower neural thresholds to sounds and better representation of higher frequencies following an enriched versus control environment. Importantly, neural gap detection improved in both quiet and background noise, indicating increased temporal processing acuity after this novel AAE intervention. Together these findings suggest that early exposure to temporally-modulated broadband noise stimuli can restrict the negative consequences of SNHL on peripheral function, and spectral and static temporal processing in the CAS.

Sensorineural damage in the cochlea decreases sensitivity and distorts auditory input from the periphery ^41^. Consequent to this peripheral damage, a number of structural and neurophysiological changes occur in brainstem, midbrain, and cortical auditory brain regions ^42-51^. Altered central auditory processing associated with SNHL, such high thresholds, loss of frequency representation, and broader tuning curves^26, 52^, may impair auditory signal detection and differentiation. Additionally, very early SNHL, experienced in DBA mice and infants with congenital SNHL, may interact with normal developmental timelines for central auditory processing to further impair sound processing beyond the direct consequences of SNHL. In normal hearing humans and rodents, temporal processing acuity, as measured by gap thresholds, improves during early development ^53, 54^. Here we find deficits in neural correlates of gap detection in the auditory midbrain in 1-month old DBA mice relative to normal hearing strains ^17, 37^. Thus, while SNHL in adulthood does not have profound effects on temporal processing in the auditory midbrain ^37^, the current findings suggest that SNHL during development may strongly impact both spectral and temporal aspects of central auditory processing.

Previous work indicates that in young mice with progressive, congenital SNHL, early exposure to broadband AAE can preserve hearing sensitivity and limit hair cell loss ^26-28, 30, 55^. This also appears to limit concomitant reorganization in the CAS, limiting the loss of neurons in the cochlear nucleus and sensitivity to high frequency sounds in the IC ^28, 30^. Here we expand this characterization of central auditory processing after early temporally modulated broadband AAE exposure, finding that intervention can preserve both spectral and temporal acuity in the auditory midbrain. After AAE intervention with our novel temporally complex AAE, and to a lesser extent with a less complex broadband AAE, neural sound processing in the IC exhibited lower response thresholds, greater high frequency encoding, more narrow frequency response areas, and better gap encoding and detection. Others have reported similar CAS plasticity following more general environmental enrichment with an auditory component. Recordings in the auditory cortex (AC) showed improved neural temporal response properties, increased spectral and temporal selectivity, and more narrow neural response fields ^56, 57^. The current findings continue to support the idea that both peripheral and central auditory processing can be modulated by enriched environments.

The plasticity of the CAS is remarkable across mammalian species. Like rodents, humans exposed to various types of passive AAE that alter sound input to the ear also undergo profound central auditory and perceptual changes resulting from neural plasticity. The effects of AAE on central auditory function, particularly in the face of hearing loss, may arise, at least in part, from homeostatic mechanisms that maintain neural activity (Turrigiano, 1999). Consistent with this idea, when sound input is attenuated via deprivation (i.e., via temporary earplug, or conductive hearing loss), the reduced peripheral input leads to increased central activity (see ^58^). Subsequent (or concurrent) exposure to passive AAE is predicted to stabilize peripheral excitatory drive and preserve input to the CAS. By stabilizing the mean level of neural activity, AAE is predicted to counteract hearing loss-related increases in central gain, improving coding efficiency, and maintaining an optimal input-dependent dynamic range ^59^. Indeed, perceptual changes in humans are observed following AAE that are consistent with normalized gain, including altered loudness perception ^60, 61^, and finer intensity resolution ^62^. AAE may also help to maintain or expand sound representation in the face of deteriorating peripheral input through standard experience-dependent plasticity mechanisms. Humans show improved temporal coding following AAE ^63^, which could arise from improved sound representation. In rodents, AAE can lead to reorganization in primary and non-primary auditory cortex as reflected in narrower response fields, improved temporal response properties, and increased spectral and temporal selectivity of neurons ^57, 64^.

Intriguingly, while both types of passive AAE employed in this study improved auditory sensitivity and spectral sound processing, our novel, temporally complex broadband AAE had a stronger positive influence on temporal processing acuity. This was particularly true for improvements in temporal processing with background noise. Thus, the benefits of early AAE exposure may be related to characteristics of the sound presented. Previous work showed that in young DBA mice, treatment with broadband AAE improved behavioral and neurophysiological measures of tonal thresholds ^26^. In the present study, a similar, but temporally more complex, broadband AAE stimulus, improved both spectral and temporal sound processing. Band-limited AAE also slowed the progression of SNHL in the 16 to 32 kHz range, but did not ameliorate a loss of sensitivity at lower frequencies ^55^. Likewise, the effect of AAE exposure is also shaped by auditory function and timing of the intervention. In mature auditory systems, or with normal hearing, some types of AAE exposure may instead lead to the suppression of sound sensitivity ^30, 65^. Recently it was observed that young adult CBA mice exposed to 75 dB SPL AAE were found to display functional evidence of cochlear synaptopathy ^66^. Likewise, in adult cats with normal hearing, tonal or band-limited AAE exposure profoundly suppressed AC activity in the frequency range of the exposure ^67-70^.

Though the precise mechanism is unknown, early broadband AAE has a positive impact on cochlear health across a limited tonotopic range depending on the spectral composition of the AAE ^28^. Improved cochlear function in turn supports lower thresholds and maintains frequency representation in the CAS. However, this study re-affirms that broadband AAE exposure invokes further adaptive neural plasticity in the auditory midbrain. Sharper IC neural tuning curves indicate that lateral inhibition is enhanced in the CAS following auditory enrichment. Moreover, temporally complex AAE may not only strengthen inhibition, but also improve the timing of inhibition. Timed inhibition is key for shaping sound offset responses that subserve gap detection ^71-74^. Since more spectrally complex signals evoke stronger sound offset responses, the broadband nature of the stimulus might be important, and may even be improved upon with a variable spectral component ^75^. The possibility that temporal encoding can be altered by auditory experience was first confirmed by Kilgard and Merzenich ^43^, who demonstrated that the ability of AC neurons to follow high frequency sound stimuli can be improved if high frequency sound stimuli are paired with electrical stimulation of the nucleus basalis. The improvement in neural encoding and detection of gaps in noise in the current study shows that even passive sound exposure may shape temporal acuity, though it seems unlikely that this occurs through pathways involving the nucleus basalis.

Overall, the current findings suggest that temporally-complex AAE interventions may provide functional benefits in individuals with SNHL, especially newborns diagnosed with hearing loss. The improved neural encoding of short gap durations in the IC is likely to support functional improvements in gap detection, as these measures are strongly associated ^17^. In turn, improved gap detection, particularly in noise, counteracts an aspect of hidden hearing loss that impairs speech perception in daily life ^12^. Though not specific to the temporally complex AAE intervention, better frequency representation and sharper spectral tuning that occurs after AAE may also bolster auditory signal detection and differentiation. These findings support the possibility that AAE may be targeted, based on the properties of the AAE as well as the listener, to better improve hearing deficits. The ability of our novel, temporally complex broadband AAE exposure to improve neural correlates of SNHL provides direct bench-to-bedside promise for treating congenital SNHL.

